# Bacterial catabolic system of acetovanillone and acetosyringone useful for upgrading aromatic compounds obtained through chemical lignin depolymerization

**DOI:** 10.1101/2022.04.28.489975

**Authors:** Yudai Higuchi, Naofumi Kamimura, Hiroki Takenami, Yusei Kikuiri, Chieko Yasuta, Kenta Tanatani, Toru Shobuda, Yuichiro Otsuka, Masaya Nakamura, Tomonori Sonoki, Eiji Masai

## Abstract

Acetovanillone is a major aromatic monomer produced in oxidative/base-catalyzed lignin depolymerization. However, the production of chemical products from acetovanillone has not been explored due to the lack of information on the microbial acetovanillone catabolic system. Here *acvABCDEF* was identified as specifically induced genes during the growth of *Sphingobium* sp. strain SYK-6 cells with acetovanillone and these genes were essential for SYK-6 growth on acetovanillone and acetosyringone (a syringyl-type acetophenone derivative). AcvAB and AcvF produced in *Escherichia coli* phosphorylated acetovanillone/acetosyringone and dephosphorylated the phosphorylated acetovanillone/acetosyringone, respectively. AcvCDE produced in *Sphingobium japonicum* UT26S converted the dephosphorylated phosphorylated acetovanillone/acetosyringone intermediate into vanilloyl acetic acid/3- (4-hydroxy-3,5-dimethoxyphenyl)-3-oxopropanoic acid through carboxylation. To demonstrate the feasibility of producing *cis*,*cis*-muconic acid from acetovanillone, a metabolic modification on a mutant of *Pseudomonas* sp. strain NGC7 that accumulates *cis*,*cis*-muconic acid from catechol was performed. The resulting strain expressing *vceA* and *vceB* required for converting vanilloyl acetic acid to vanillic acid and *aroY* encoding protocatechuic acid decarboxylase in addition to *acvABCDEF* successfully converted 1.2 mM acetovanillone to approximate equimolar *cis*,*cis*-muconic acid. Our results are expected to help improve the yield and purity of value-added chemical production from lignin through biological funneling.

**IMPORTANCE:** In the alkaline oxidation of lignin, aromatic aldehydes (vanillin, syringaldehyde, and *p*-hydroxybenzaldehyde), aromatic acids (vanillic acid, syringic acid, and *p*- hydroxybenzoic acid), and acetophenone-related compounds (acetovanillone, acetosyringone, and 4’-hydroxyacetophenone) are produced as major aromatic monomers. Also, base-catalyzed depolymerization of guaiacyl lignin resulted in vanillin, vanillic acid, guaiacol, and acetovanillone as primary aromatic monomers. To date, microbial catabolic systems of vanillin, vanillic acid, and guaiacol have been well characterized, and the production of value-added chemicals from them has also been explored. However, due to the lack of information on the microbial acetovanillone and acetosyringone catabolic system, chemical production from acetovanillone and acetosyringone has not been achieved. This is the first study to elucidate the acetovanillone/acetosyringone catabolic system, and to demonstrate the potential of using these genes for value-added chemicals production from these compounds.

## INTRODUCTION

The use of lignocellulosic biomass is expected to build a decarbonized society that breaks away from fossil resources. The development of lignin utilization has become particularly important in the utilization of lignocellulose. Lignin content in lignocellulose ranges from 9% to 32% (1), and gymnosperm (softwood) lignins are composed of guaiacyl (G) units with small amounts of *p*-hydroxyphenyl (H) units, whereas angiosperm (hardwood) lignins are composed of G-units and syringyl (S) units (2, 3). Grass lignins contain G- and S-units and more H-units than gymnosperm lignins (2, 3). Because of lingin’s complex and heterogeneous structure, it is difficult to selectively extract specific compounds even when it is degraded, and it has been ineffectively used so far (1, 4). Recently, “biological funneling,” in which a heterogeneous mixture of low-molecular-weight aromatic compounds obtained from chemo-catalytic depolymerization of lignin is converged to specific value-added chemicals, such as *cis*,*cis*-muconic acid (ccMA) and 2-pyrone-4,6-dicarboxylic acid using microbial catabolic functions, has attracted attention (5–12).

In the alkaline oxidation of lignin, aromatic aldehydes (vanillin, syringaldehyde, and *p*-hydroxybenzaldehyde), aromatic acids (vanillic acid, syringic acid, and *p*- hydroxybenzoic acid), and acetophenone-related compounds [acetovanillone (AV), acetosyringone (AS; a syringyl-type acetophenone derivative), and 4’- hydroxyacetophenone (a *p*-hydroxyphenyl-type acetophenone derivative)] are produced as major aromatic monomers (13–15). Base-catalyzed Indulin AT (a pine kraft lignin) depolymerization resulted in the formation of vanillin, vanillic acid, guaiacol, and AV as major aromatic monomers, similar to the alkaline oxidation of Lignoboost lignin (a softwood kraft technical lignin) (8, 16–19). A black liquor produced in a softwood kraft pulping process, Lignoforce, contained guaiacol, vanillin, and AV as major aromatic monomers (20). Among these major aromatic monomers, microbial catabolic systems of vanillin, vanillic acid, and guaiacol have been characterized (21–29). Additionally, value-added chemical production from vanillin, vanillic acid, and guaiacol has also been explored (12, 18, 30–36). However, due to the lack of information on the microbial AV catabolic system, chemical production from AV has not been reported (16, 18, 19).

Therefore, chemical production from all of the major aromatic monomers produced by oxidative/base-catalyzed lignin depolymerization has been unachieved. Recently, Eltis and coworkers isolated *Rhodococcus rhodochrous* GD01 and GD02, which can utilize AV in addition to vanillin, vanillic acid, and guaiacol in the black liquor produced during the Lignoforce kraft pulping of softwood (20). Based on the observation that *apkC* was induced when GD02 was cultured in black liquor extracts containing AV, *apkA*-*apkB*-*apkC*, presumed to encode biotin-dependent carboxylase, were predicted to be involved in AV catabolism. However, the enzymatic system encoded by these genes is unknown. Additionally, in *Arthrobacter* sp. strain TGJ4, 4’-hydroxyacetophenone and AV are converted to 4-hydroxybenzoic acid and vanillic acid, respectively; however, the enzymatic system has been unclarified (37).

*Sphingobium* sp. strain SYK-6 can utilize various lignin-derived dimers, such as β- aryl ether, biphenyl, phenylcoumaran, and diarylpropane, as well as monomers, including ferulic acid, vanillin, vanillic acid, and AV (11, 16, 20, 38, 39). In SYK-6 cells, stereoisomers of β-aryl ether model compound, guaiacylglycerol-β-guaiacyl ether, are stereospecifically converted to achiral β-hydroxypropiovanillone (HPV) via Cα- oxidation, ether cleavage, and glutathione removal (Fig. 1) (38, 40–44). HPV and β- hydroxypropiosyringone (HPS, a syringyl-type β-aryl ether metabolite) are oxidized to vanilloyl acetic acid (VAA) and 3-(4-hydroxy-3,5-dimethoxyphenyl)-3-oxopropanoic acid (SAA), respectively, and further catabolized via vanillic acid and syringic acid. (Fig. 1) (45). Additionally, we identified *vceA* and *vceB*, which encode an acetyl-CoA-dependent VAA/SAA-converting enzyme and a vanilloyl-CoA/syringoyl-CoA thioesterase, respectively (5). However, these enzyme genes were found to be not essential for VAA/SAA catabolism in SYK-6. It has been reported that VAA is chemically unstable and can be converted to AV by a non-enzymatic decarboxylation (46, 47). Indeed, when an SYK-6 cell extract was incubated with HPV, a small amount of AV along with VAA was generated (Fig. 1) (45).

**FIG 1.**
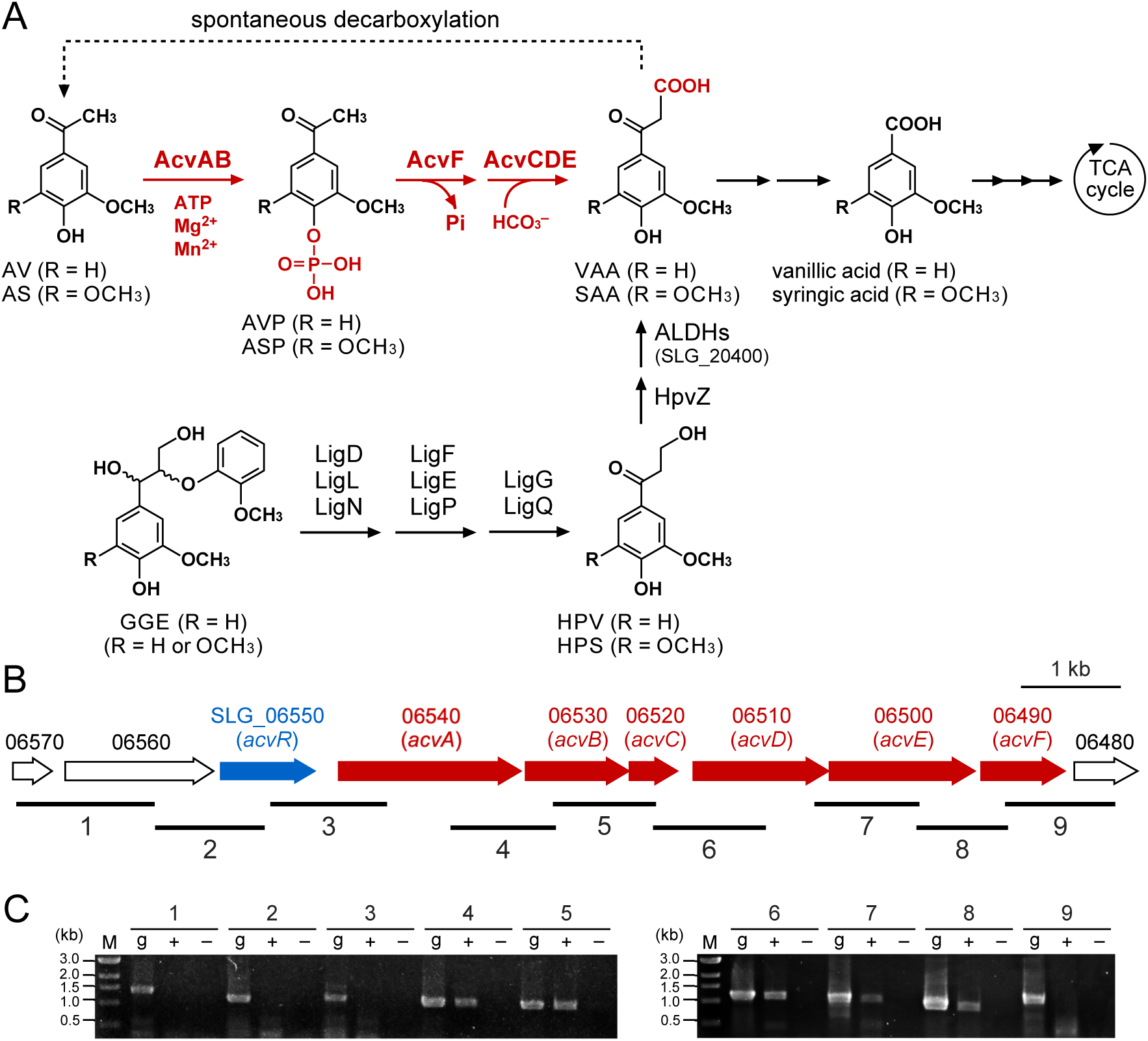
(A) Catabolic pathway of AV and AS in *Sphingobium* sp. strain SYK-6. The pathways for both guaiacyl (R = H)- and syringyl (R = OCH_3_)-type compounds are shown. VAA, an intermediate metabolite of GGE, has been suggested to be spontaneously decarboxylated to AV (Higuchi et al., 2018; Niwa and Saburi, 2002). Enzymes: AcvAB, AVP/ASP synthetase; AcvF, AVP/ASP phosphatase; AcvCDE, biotin-dependent carboxylase; LigD, LigL, and LigN, Cα-dehydrogenases; LigF, LigE, and LigP, β-etherases; LigG and LigQ, glutathione *S*-transferases; HpvZ, HPV/HPS oxidase; ALDHs, aldehyde dehydrogenases; SLG_20400, vanilloyl acetaldehyde dehydrogenase. Abbreviations: AV, acetovanillone; AS, acetosyringone; AVP, 4-acetyl-2-methoxyphenylphosphate; ASP, 4-acetyl-2,6- dimethoxyphenylphosphate; VAA, vanilloyl acetic acid; SAA, 3-(4-hydroxy-3,5-dimethoxyphenyl)-3- oxopropanoic acid; GGE, guaiacylglycerol-β-guaiacyl ether; HPV, β-hydroxypropiovanillone; HPS, β- hydroxypropiosyringone. (B) Gene organization of *acvABCDEF*. Arrows indicate the genes from SLG_06570 to SLG_06480. (C) RT-PCR analysis of *acvABCDEF*. Total RNA used for cDNA synthesis was isolated from SYK-6 cells grown in Wx-SEMP containing 5 mM AV. The regions to be amplified are indicated by black bars below the genetic map. Lanes: M, molecular size markers; g, control PCR with the SYK-6 genomic DNA; ‘+’ and ‘−’, RT-PCR with and without reverse transcriptase, respectively.

In this study, the catabolic pathway of AV and AS in SYK-6 was determined, an SYK-6 gene cluster consisting of six novel genes involved in AV and AS catabolism was identified, and their functions clarified (Fig. 1). Furthermore, these genes were expressed in a derivative strain of *Pseudomonas* sp. strain NGC7 (Shinoda et al., 2019), which is expected to be a chassis microorganism for converting lignin-related compounds, and this engineered strain successfully produced ccMA from AV.

## RESULTS

### Determination of the AV catabolism pathway in *Sphingobium* sp. SYK-6

Intermediate metabolites generated during SYK-6 incubation with AV were identified to determine the catabolic pathway of AV in SYK-6. SYK-6 cells grown with AV were incubated with 1 mM AV in Wx minimal medium (48) for 33 h, and high-performance liquid chromatography (HPLC)–mass spectrometry (MS) analyzed the supernatant of the reaction mixture. This analysis indicated that AV disappeared and was converted into compounds I and II with retention times of 2.2 and 2.5 min, respectively (Fig. 2A and B). Based on the comparison of the retention time and the mass spectrum of compound I with those of the authentic sample, this compound was identified as vanillic acid (molecular weight [MW], 168) (Fig. 2C and E, Fig. S1). Furthermore, compound II was found to be VAA (MW, 210) by comparing with the retention time and the mass spectrum of the authentic sample (Fig. 2D and Fig. S1). These results suggest that AV was carboxylated, converted to VAA, and catabolized via vanillic acid in SYK-6 (Fig. 1).

**FIG 2.**
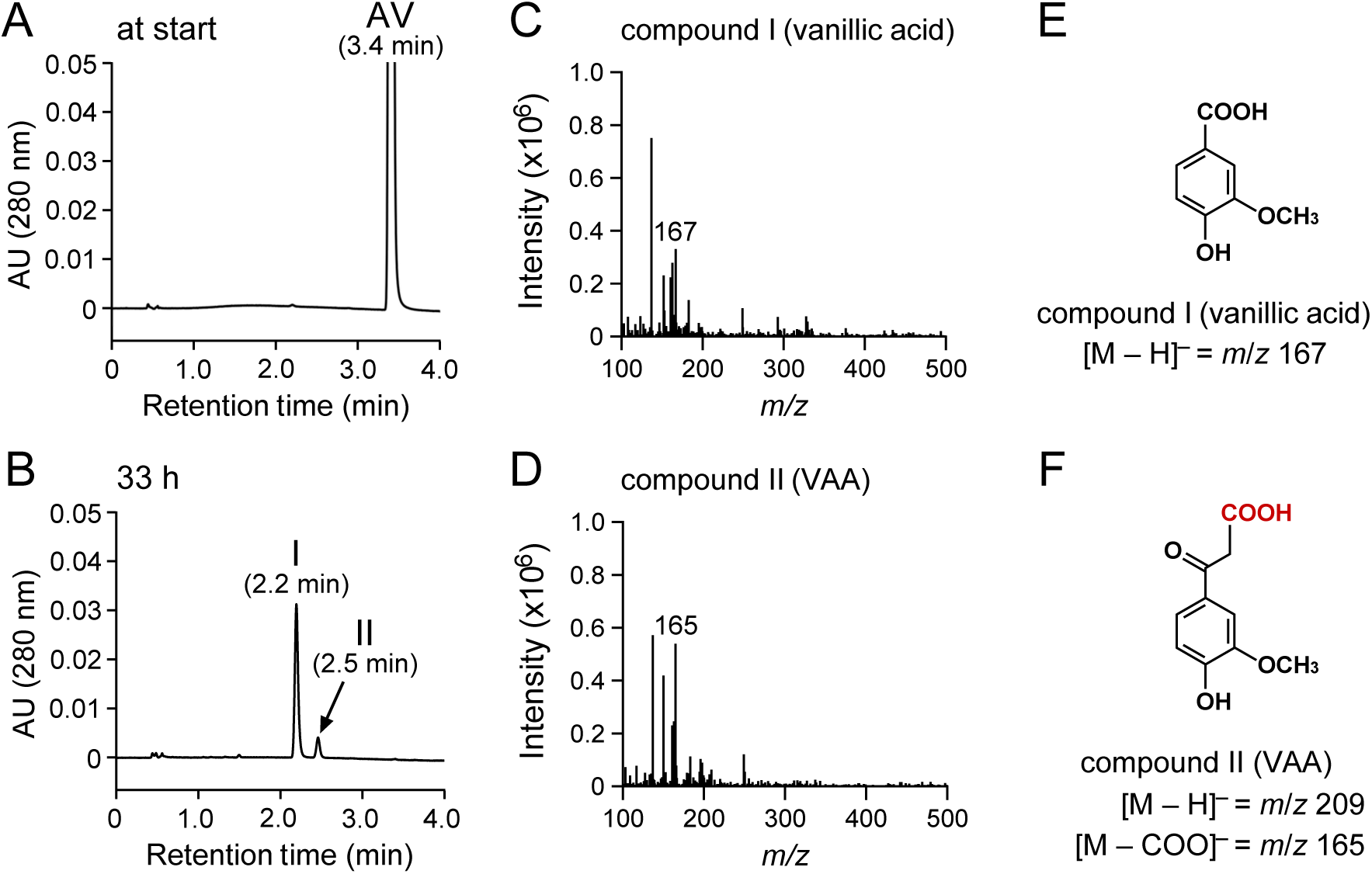
HPLC–MS analysis of AV metabolites. Cells of SYK-6 grown with AV (OD_600_ = 0.2) were incubated with 1 mM AV in Wx medium. Portions of the reaction mixtures were collected at the start (A) and after 33 h (B) of incubation and then analyzed by HPLC–MS. The ESI-MS spectra of compounds I and II (negative mode) are shown in panels C and D, respectively. (E and F) Chemical structures of compound I (vanillic acid) and compound II (VAA), respectively.

### Search for genes involved in the conversion of AV using microarray analysis

The AV conversion rate was measured using SYK-6 cells grown in Wx-containing 10 mM sucrose, 10 mM glutamic acid, 0.13 mM methionine, and 10 mM proline (Wx-SEMP; the non-inducing condition) and Wx-SEMP containing 5 mM AV (the inducing condition) to investigate the inducibility of converting AV in SYK-6. The conversion rate of 1 mM AV by cells grown under the inducing condition was approximately 27-times higher than that of cells grown under the non-inducing condition (Fig. S2). Thus, the AV-converting enzyme gene(s) was induced by AV and/or its intermediates.

The AV-converting enzyme gene(s) was searched based on the induction profiles of the entire SYK-6 genes analyzed by previous DNA microarray analysis (49). This analysis indicated that six consecutive genes consisting of SLG_06540, SLG_06530, SLG_06520, SLG_06510, SLG_06500, and SLG_06490 were induced 17–76-fold, during SYK-6 growth with AV (Fig. 1B, Table S1). Among these genes, SLG_06540 and SLG_06530 showed 44% and 41% amino acid sequence identity, respectively, with Proteins 1 and 2, which comprise phenylphosphate synthase involved in biotin and thiamine diphosphate-independent phenol carboxylation by *Thauera aromatica* K172 (Table S2) (50–52). SLG_06520 and SLG_06510 showed 36% and 50% amino acid sequence identity with a biotin carboxyl carrier protein (BCCP; AccB) and biotin carboxylase (BC; AccC) of biotin-dependent acetyl-CoA carboxylase of *Bacillus subtilis* 168, respectively (Table S2) (53). SLG_06500 showed 26% amino acid sequence identity with the carboxytransferase (CT; PycB) of biotin-dependent pyruvic acid carboxylase of *Methanothermobacter thermautotrophicus* ΔH (Table S2) (54).

These facts suggest that SLG_06520, SLG_06510, and SLG_06500 encode BCCP, BC, and CT, of biotin-dependent carboxylase, respectively (Tong, 2013). The proteomic analysis of the *Aromatoleum aromaticum* EbN1 grown with 4’-hydroxyacetophenone revealed the upregulation of XccA and XccB, presumably biotin-dependent carboxylase components, and *xccA*-*xccC*-*xccB* are predicted to be involved in 4’- hydroxyacetophenone carboxylation (55). XccA (CT), XccC (BC), and XccB (BCCP) have 55%, 63%, and 45% amino acid sequence identity with SLG_06500, SLG_06510, and SLG_06520, respectively (Table S2). Furthermore, SLG_06500, SLG_06510, and SLG_06520 exhibited 42%, 51%, and 38% amino acid sequence identity, respectively, with *apkA* (CT), *apkB* (BC), and *apkC* (BCCP), the recently predicted AV catabolic enzyme genes in *R. rhodochrous* GD02 (Table S2) (20). SLG_06490 showed 23% amino acid sequence identity with NagD, a ribonucleotide monophosphatase of *E. coli* K-12, belonging to the haloacid dehydrogenase (HAD) superfamily (Table S2) (56).

In *T. aromatica* K172, phenol is phosphorylated to phenylphosphate by phenylphosphate synthase (Proteins 1 and 2) before the carboxylation, facilitated by an accessory protein (Protein 3) (50–52). Subsequently, the core enzyme composed of α, β, and γ components catalyzing the carboxylation using CO_2_ as a substrate and the 8 subunit catalyzing the dephosphorylation are involved in converting phenylphosphate into 4-hydroxybenzoic acid via phenolate anion (57–59). Based on the above, AV carboxylation in SYK-6 was expected to proceed as follows. i) Putative 4-acetyl-2- methoxyphenylphosphate (AVP) synthetase encoded by SLG_06540 and SLG_06530 phosphorylates AV. ii) Putative phosphatase encoded by SLG_06490 dephosphorylates the phosphorylated AV. iii) Putative biotin-dependent carboxylase encoded by SLG_06520, SLG_06510, and SLG_06500 carboxylates the AV dephosphorylated anion intermediate.

Reverse transcription (RT)-polymerase chain reaction (PCR) analysis was conducted using total RNA prepared from the SYK-6 cells grown in the presence of AV to examine whether SLG_06540–SLG_06490 form a transcription unit. Specific amplification was observed from SLG_06540 to SLG_06490, indicating that SLG_06540−SLG_06490 form an operon (Fig. 1B and C).

#### Roles of SLG_06540–SLG_06490 in AV and AS catabolism

To examine whether SLG_06540−SLG_06490 are involved in AV catabolism in SYK-6, disruption mutants of SLG_06540−SLG_06490 (ι106540–ι106490) were created via homologous recombination (Fig. S3). The growth of ι106540–ι106490 cells on AV was evaluated.

Since SYK-6 cannot grow at a concentration of several millimolar of AV, 1 mM AV was added to the Wx medium at the beginning of cultivation, and another 1 mM AV was added after 52 h of incubation. The OD_660_ of SYK-6 increased after 20 h of cultivation, and further growth was observed when AV was added after 52 h. In contrast, all mutants completely lost their capacity to grow on AV (Fig. S4A). Additionally, each mutant also completely lost the capacity to grow on AS (Fig. S4B). These results indicate that SLG_06540–SLG_06490 are essential for AV and AS catabolism; thus, we designated these genes *acvA–acvF*.

#### SLG_06550 is the transcriptional regulator of *acvABCDEF*

SLG_06550, located just upstream of *acvA*, showed 25% amino acid sequence identity with NphR, an AraC-type transcriptional regulator that positively regulates the 4-nitrophenol monooxygenase gene (*nphA1A2*) of *Rhodococcus* sp. strain PN1 (60). To examine whether SLG_06550 is involved in the transcriptional *acvABCDEF* regulation in SYK- 6, an SLG_06550 disruption mutant (ι106550) was created and its capacity to grow on Wx medium containing AV was examined (Fig. S5A and B). Δ06550 completely lost its capacity to grow on AV (Fig. S5C), suggesting that SLG_06550 positively regulates the *acvABCDEF* operon, and this gene was named *acvR*.

#### *acvABCDEF* confers a host strain the capacity to carboxylate AV and AS

To examine whether *acvABCDEF* encodes the AV and AS carboxylase system, a plasmid carrying *acvABCDEF* in pJB861 was introduced into a host strain, *Sphingobium japonicum* UT26S, which is incapable of AV and AS conversion. The UT26S genome contains genes that show 32% (SJA_C1-33200) and 50% (SJA_C1-33210) amino acid sequence identity with *acvC* and *acvD,* encoding putative BCCP and BC, respectively, but no ortholog of *acvA* and *acvB* (encoding putative AVP synthetase), *acvE* (encoding putative CT), and *acvF* (encoding putative phosphatase). The resting cells of UT26S expressing *acvABCDEF* [optical density at 600 nm (OD_600_) = 10.0] were incubated with 200 µM AV for 1 h. HPLC–MS analysis showed that VAA was produced (Fig. 3A–D). When the same resting cells were incubated with 200 µM AS for 4 h, compound III with a retention time of 3.4 min was produced (Fig. 3E and F). Negative electrospray ionization-MS (ESI-MS) analysis of compound III showed a major fragment at *m/z* 239 (Fig. 3G and H), suggesting that compound III was SAA (MW, 240). These results indicate that *acvABCDEF* encodes components of the carboxylase system required for AV and AS catabolism.

**FIG 3.**
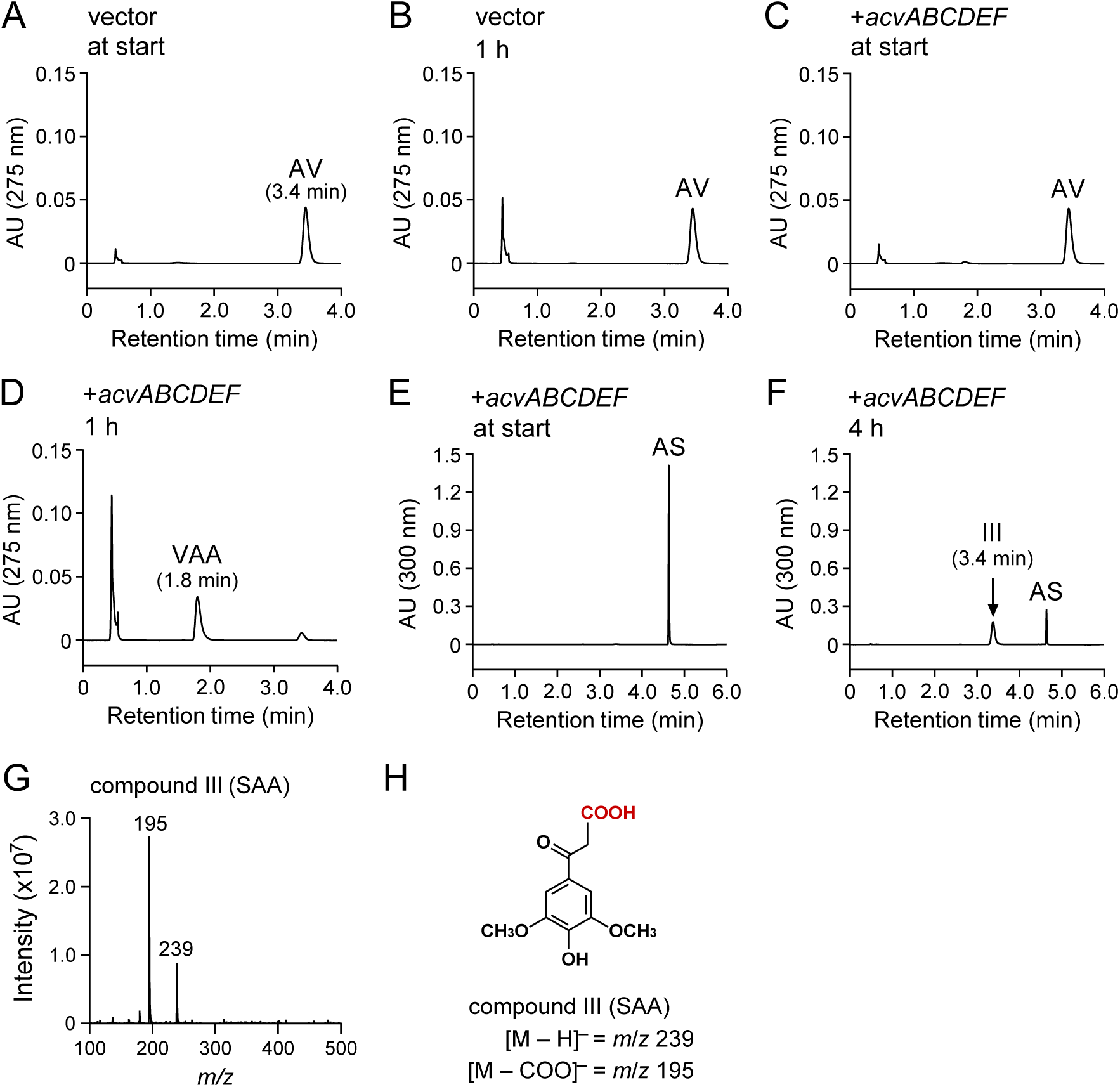
Conversions of AV and AS by resting cells of *S. japonicum* UT26S carrying *acvABCDEF*. Resting cells of UT26S harboring pJB861 (OD_600_ = 10.0; A and B) and resting cells of UT26S harboring pJB*acv* (OD_600_ = 10.0; C–F) were incubated with AV (200 µM; A–D) or AS (200 µM; E and F). Portions of the reaction mixtures were collected at the start (A, C, and E), after 1 h (B and D), and after 4 h (F) of incubation and analyzed by HPLC–MS. The ESI-MS spectrum of compound III (negative mode) is shown in panel G. (H) Chemical structure of compound III (SAA).

### A mixture of AcvA and AcvB catalyzes the phosphorylation of AV and AS

Proteins 1 and 2 (phenylphosphate synthase) of *T. aromatica* K172, where *acvA* and *acvB,* respectively, show amino acid sequence similarity, convert phenol to phenylphosphate by the coexistence of both (51). MgATP is essential as a phosphoryl donor for this conversion and Mn^2+^ promotes catalytic activity (51). Each *acvA* and *acvB* fused with a His-tag at the 5’ terminus was expressed in *E. coli* to characterize the function of *acvA* and *acvB.* Sodium dodecyl sulfate-polyacrylamide gel electrophoresis (SDS-PAGE) analysis showed the production of 71-kDa and 40-kDa proteins in cell extracts of *E. coli* expressing His-tagged *acvA* and *acvB,* respectively (Fig. S6).

Crude AcvA (500 µg/mL), AcvB (500 µg/mL), and AcvA + AcvB (500 µg/mL each) reacted with 200 µM AV, respectively, in the presence of 2 mM ATP, 2 mM MgCl_2_, and 200 μM MnCl_2_. HPLC–MS analysis showed that AV was completely converted into compound IV (1.9 min) after 30 min of incubation when AcvA and AcvB coexisted (Fig. 4A–C). Negative ESI-MS analysis of compound IV showed a fragment at *m*/*z* 245. Compound IV was identified as AVP (MW, 246) based on the molecular weight deduced from the fragment ion (Fig. 4D and E). Therefore, it was suggested that AcvAB phosphorylated the hydroxy group of AV.

**FIG 4.**
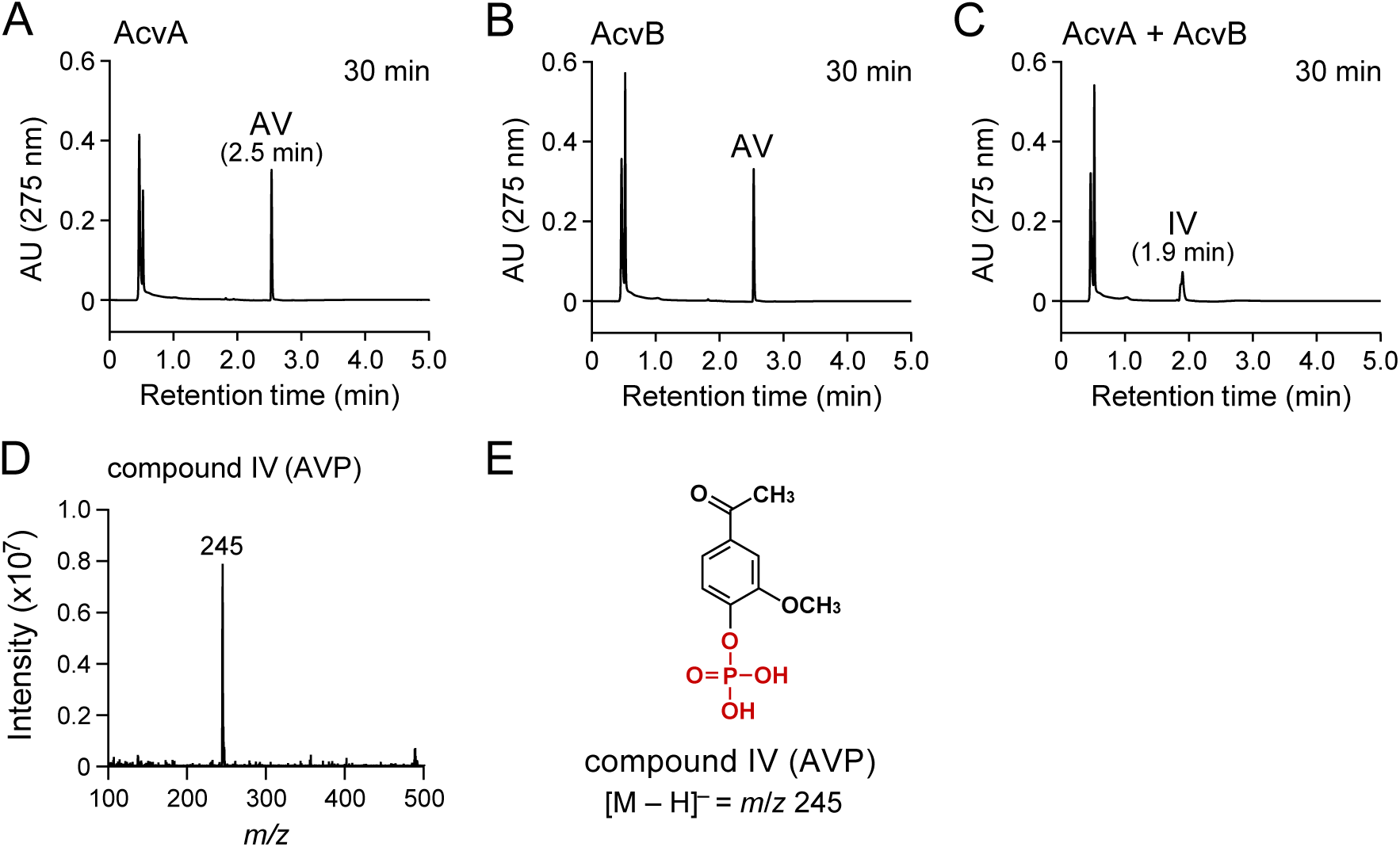
Conversion of AV by crude AcvA and AcvB. AV (200 µM) was incubated with a mixture of cell extracts of *E. coli* BL21(DE3) harboring pE16*acvA* and *E. coli* BL21(DE3) harboring pET-16b (500 µg protein/mL each; A), a mixture of cell extracts of *E. coli* BL21(DE3) harboring pE16*acvB* and *E. coli* BL21(DE3) harboring pET-16b (500 µg protein/mL each; B), and a mixture of cell extracts of *E. coli* BL21(DE3) harboring pE16*acvA* and *E. coli* BL21(DE3) harboring pE16*acvB* (500 µg protein/mL each; C). Reactions were performed in the presence of 2 mM ATP, 2 mM MgCl_2_, and 200 µM MnCl_2_. Portions of the reaction mixtures were collected after 30 min of incubation and analyzed by HPLC. The ESI-MS spectrum of compound IV (negative mode) is shown in panel D. (E) Chemical structure of compound IV (AVP).

AcvA and AcvB were purified to near homogeneity by Ni affinity chromatography from the cell extracts of *E. coli* expressing His-tagged *acvA* and *acvB*, respectively (Fig. S6). However, because the specific activity of purified AcvA + AcvB was lower than that of crude AcvA + AcvB, the AcvAB enzyme properties were investigated using crude enzymes.

To examine the AcvAB cofactor requirement, crude AcvA + AcvB (50–1000 μg protein/mL each) was incubated with 100 µM AV in the presence and absence of cofactors (2 mM ATP, 2 mM MgCl_2_, 200 µM MnCl_2_, 2 mM ATP + 2 mM MgCl_2_, 2 mM ATP + 200 µM MnCl_2_, 2 mM MgCl_2_ + 200 µM MnCl_2_, and 2 mM ATP + 2 mM MgCl_2_ + 200 µM MnCl_2_). The highest activity (ca. 111 nmol·min^−1^·mg^−1^) was obtained in the presence of ATP + Mg^2+^ + Mn^2+^, whereas 42% and 3% activities were observed with ATP + Mg^2+^ and ATP + Mn^2+^, respectively. No activity was observed in the presence of the other cofactors. These results indicate that AcvAB used MgATP as a phosphoryl donor for AV phosphorylation similar to the phenylphosphate synthase of *T. aromatica* K172, and that phosphorylation activity is promoted in the presence of Mn^2+^.

Crude AcvA + AcvB (10–500 μg protein/mL each) was incubated with 100 μM AV, AS, acetophenone, 4’-hydroxyacetophenone, 3’-hydroxyacetophenone, 3’,4’- dihydroxyacetophenone, 3’-hydroxy-4’-methoxyacetophenone, 3’,4’- dimethoxyacetophenone, 3’,4’,5’-trimethoxyacetophenone, 4’-hydroxypropiophenone, 4’-hydroxybuthyrophenone, 4’-hydroxyvalerophenone, guaiacol, vanillic acid, and 4- hydroxybenzoic acid, respectively, in the presence of 2 mM ATP + 2 mM MgCl_2_ + 200 μM MnCl_2_ to examine the substrate range of AcvAB (Fig. S7). HPLC–MS analysis showed that AcvAB converted AS into 4-acetyl-2,6-dimethoxyphenylphosphate (ASP) (Fig. S8A–C). AcvAB exhibited activity toward compounds with the hydroxy group at the 4-position of the aromatic ring except for vanillic acid and 4-hydroxybenzoic acid (Fig. S8). AcvAB also showed activity toward 3’-hydroxyacetophenone but it showed no activity toward 3’-hydroxy-4’-methoxyacetophenone.

#### AcvF catalyzes the dephosphorylation of AVP and ASP

Because AcvF showed 23% amino acid sequence identity with NagD of *E. coli* K-12, belonging to the HAD superfamily, including phosphatases, AcvF is expected to have dephosphorylation activities for AVP and ASP. (56). The AcvF function was characterized by expressing *acvF* fused with a His-tag at the 5’ terminus in *E. coli*. SDS-PAGE analysis showed a 32-kDa protein production in cell extract of *E. coli* expressing His-tagged *acvF*, and AcvF was purified to near homogeneity by Ni affinity chromatography (Fig. S6). To investigate the AVP and ASP dephosphorylation ability of AcvF, AVP and ASP were produced from AV and AS, respectively, using crude AcvA + AcvB. Crude AcvA + AcvB (1 mg protein/mL each) was incubated with 200 μM AV and AS, respectively, in the presence of 2 mM ATP + 2 mM MgCl_2_ + 200 μM MnCl_2_ for 2 h. HPLC analysis confirmed that AV and AS were completely converted into AVP and ASP, respectively, and the filtrate obtained via ultrafiltration (MW cutoff, 10 kDa) was used as substrates. Purified AcvF (5 µg protein/mL) was incubated with 100 μM AVP or ASP for 60 min. HPLC analysis of the reaction mixtures showed that AcvF converted AVP into AV (Fig. 5A and B). Similarly, AcvF converted ASP into AS (Fig. 5C and D). These results indicated that AcvAB phosphorylates AV and AS, which is then dephosphorylated by AcvF, respectively.

**FIG 5.**
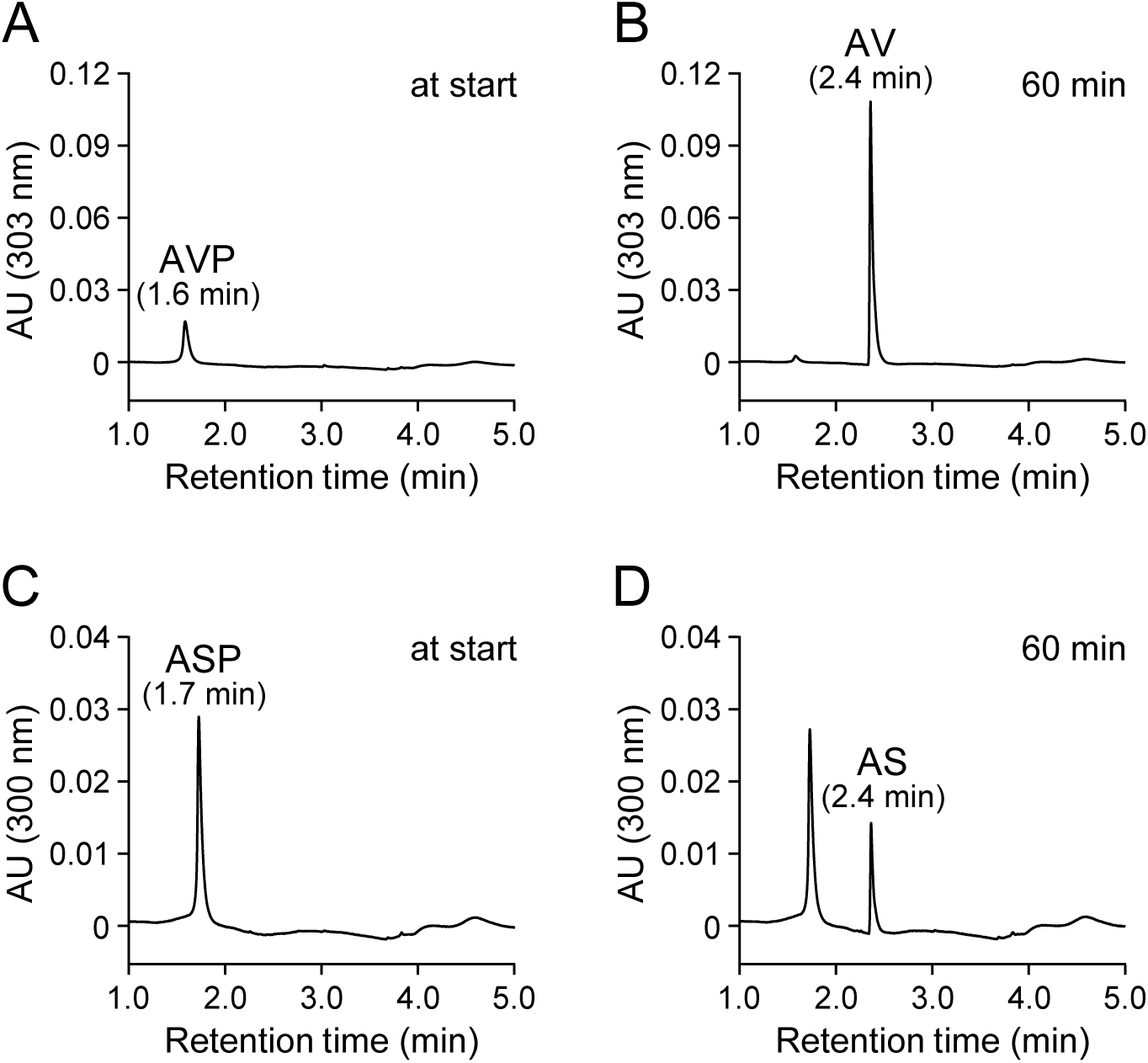
Conversions of AVP and ASP by AcvF. AVP (100 µM; A and B) and ASP (100 µM; C and D) were incubated with purified AcvF (5 µg protein/mL). Portions of the reaction mixtures were collected at the start (A and C) and 60 min (B and D) of incubation and analyzed by HPLC.

#### Carboxylation of AV and AS by a mixture of AcvAB, AcvF, and AcvCDE

It was expected that AcvC-AcvD-AcvE could carboxylate the acetyl groups of AV and AS since *acvC*, *acvD*, and *acvE* was predicted to encode biotin-dependent carboxylase components, BCCP, BC, and CT, respectively (61). A His-tag was fused to the 5’ terminus of *acvC* to express *acvCDE* in *E. coli*, characterizing the function of *acvC*, *acvD*, and *acvE.* However, SDS-PAGE analysis showed no clear AcvD and AcvE production, except for AcvC (Fig. S9). Furthermore, the resulting cell extract of *E. coli* expressing *acvCDE* (1 mg protein/mL) was incubated with 100 μM AV in the presence of crude AcvA + AcvB (1 mg protein/mL each), purified AcvF (10 μg protein/mL), 2 mM ATP + 2 mM MgCl_2_ + 200 µM MnCl_2_, and 10 mM NaHCO_3_ for 60 min; however, no reaction product was observed (data not shown). Therefore, *acvCDE* was expressed under the control of the Q5 promoter of the pQF vector using *Sphingobium japonicum* UT26S as a host. The resulting cell extract of UT26S expressing *acvCDE* (crude AcvCDE) reacted with 100 µM AV under the same reaction conditions as above. The conversion product was undetected when the cell extract of UT26S harboring pQF vector was used (Fig. 6A and B), whereas VAA was detected in the reaction with crude AcvCDE (Fig. 6C). Additionally, SAA was detected when AS was used as a substrate (Fig. 6D–F). These results suggest that AcvCDE catalyzes AV and AS carboxylation.

**FIG 6.**
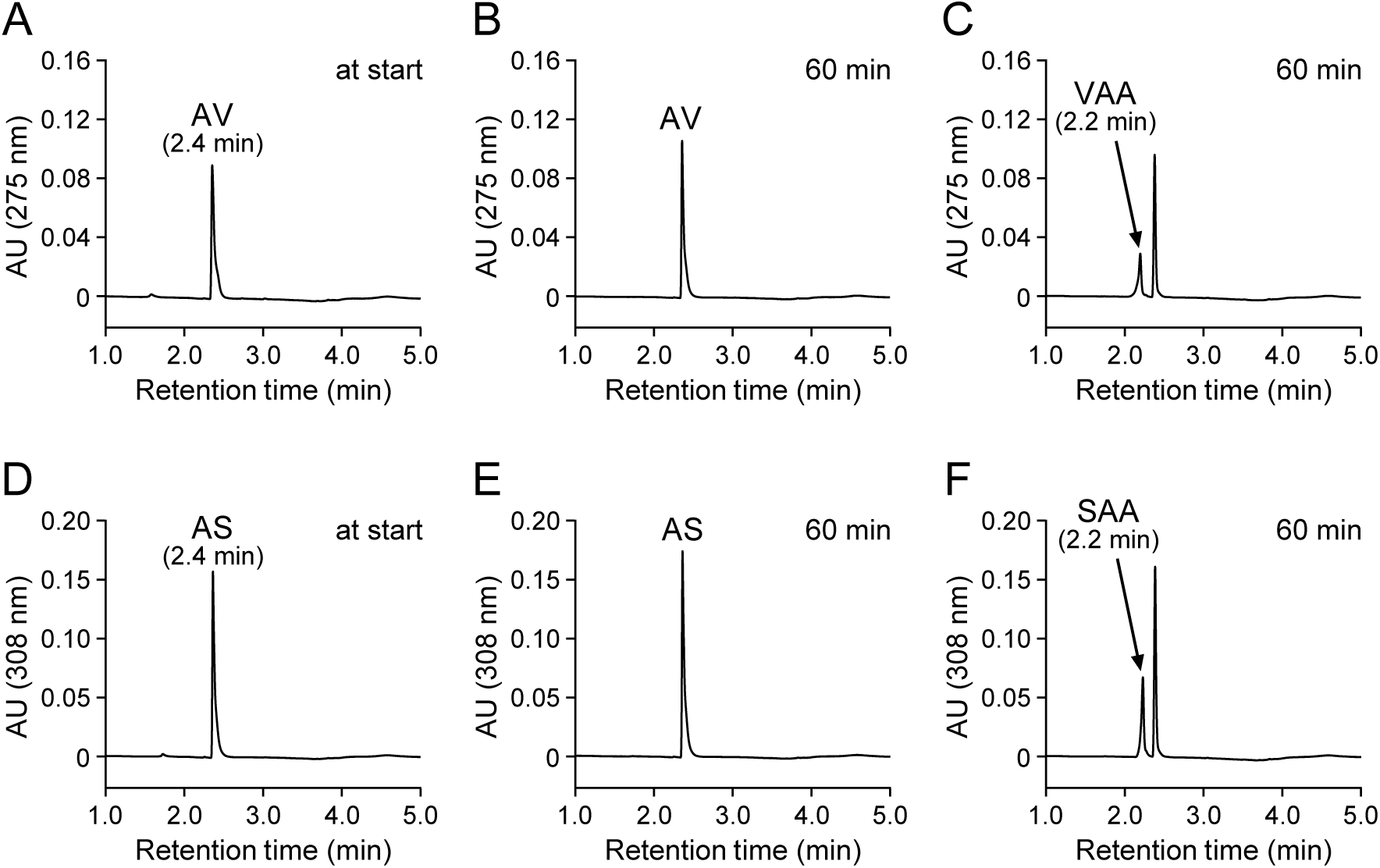
A mixture of AcvA-AcvB, AcvF, and AcvC-AcvD-AcvE catalyzed carboxylation of AV and AS. AV (100 µM; A–C) and AS (100 µM; D–F) were incubated with a cell extract of *S. japonicum* UT26S harboring pQF (1 mg protein/mL; A, B, D, and E) or a cell extract of UT26S harboring pQF*acvCDE* (1 mg protein/mL; C and F) in the presence of AcvA-AcvB and AcvF. Specifically, the reactions were performed in the presence of a cell extract of *E. coli* BL21(DE3) harboring pE16*acvA* (1 mg protein/mL), a cell extract of *E. coli* BL21(DE3) harboring pE16*acvB* (1 mg protein/mL), purified AcvF (10 µg protein/mL), 2 mM ATP, 2 mM MgCl_2_, 200 µM MnCl_2_, and 10 mM NaHCO_3_. Portions of the reaction mixtures were collected at the start (A and D) and after 60 min (B, C, E, and F) of incubation and analyzed by HPLC.

To examine whether AcvCDE directly carboxylates AV, crude AcvCDE (1 mg protein/mL) was incubated with 100 μM AV in the presence of 2 mM ATP + 2 mM MgCl_2_ + 200 μM MnCl_2_, and 10 mM NaHCO_3_ for 60 min, but no VAA formation was observed (Fig. S10A and B). Conversely, when AVP was used as a substrate, VAA was generated under the same reaction condition except for the substrate (Fig. S10C and D). These results indicate that AcvCDE could not directly carboxylate AV. Although *acvF* is essential for SYK-6 growth on AV, crude AcvCDE converted AVP into VAA in the absence of AcvF. When AVP was incubated with a cell extract of UT26S harboring pQF, conversion of AVP into AV was shown (Fig. S10E). Therefore, enzyme(s) present in UT26S appear to complement the AcvF function.

### Expression of acvABCDEF with vceA, vceB, and aroY in Pseudomonas sp

#### NGC703 enables ccMA production from AV

Our previous study demonstrated ccMA production from vanillic acid using *Pseudomonas* sp. NGC703 [a mutant of *Pseudomonas* sp. NGC7 deficient in the protocatechuic acid 3,4-dioxygenase gene (*pcaHG*) and ccMA cycloisomerase gene (*catB*)] cells harboring pTS084, comprising the protocatechuic acid decarboxylase (*aroY*), flavin prenyltransferase (*kpdB*), vanillic acid *O*-demethylase (*vanAB*), and catechol 1,2-dioxygenase (*catA*) genes (Fig. 7A) (34). This study examined whether ccMA production from AV is possible using NGC7 as a platform.

**FIG 7.**
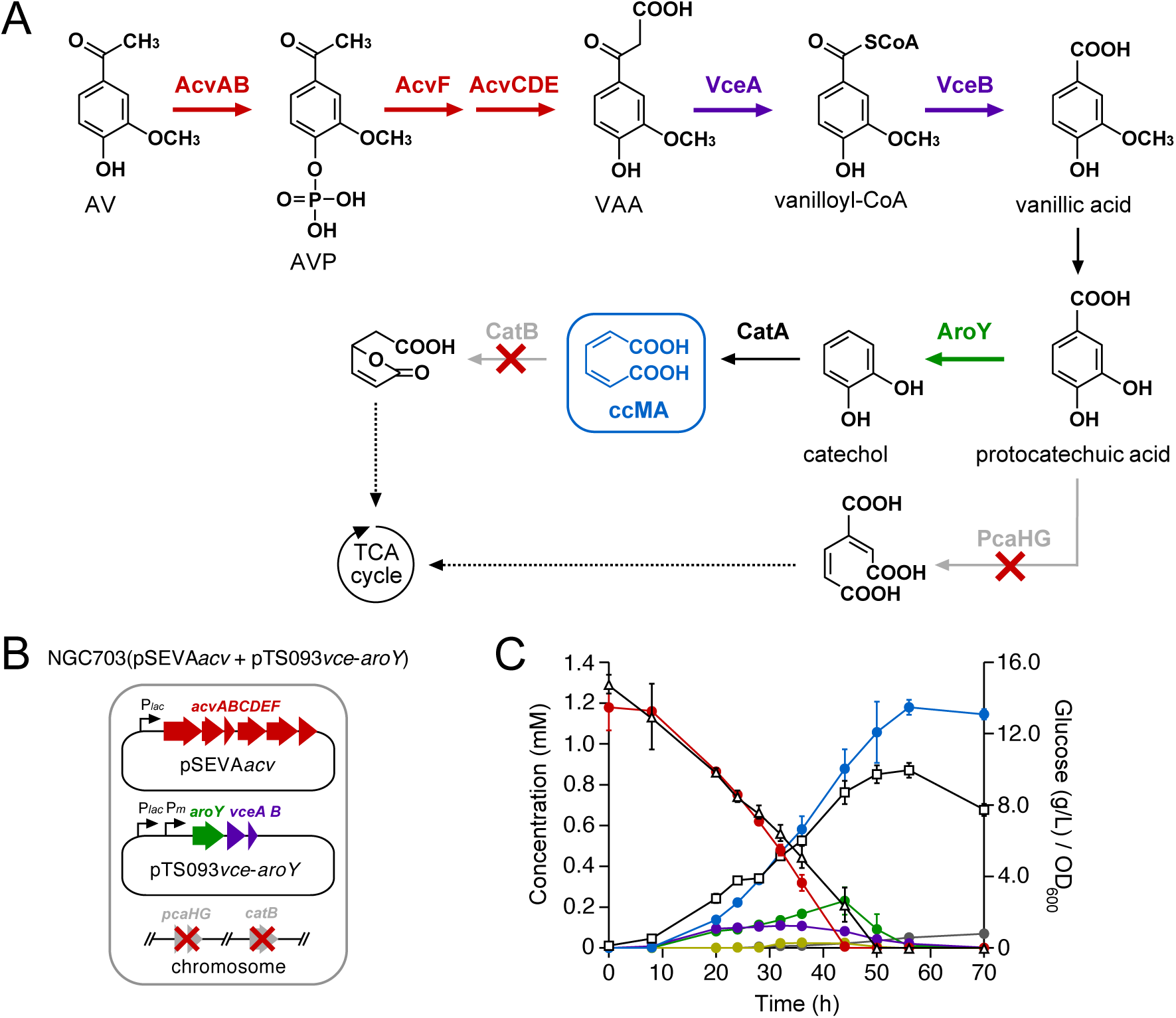
Production of ccMA from AV through the engineered metabolic pathway constructed in *Pseudomonas* sp. NGC7. (A) Engineered route for ccMA production from AV. Enzymes: AcvAB, AVP/ASP synthetase; AcvF, AVP/ASP phosphatase; AcvCDE, biotin-dependent carboxylase; VceA, VAA/SAA-converting enzyme; VceB, vanilloyl-CoA/syringoyl-CoA thioesterase; AroY, protocatechuic acid decarboxylase; PcaHG, protocatechuic acid 3,4-dioxygenase; CatA, catechol 1,2-dioxygenase; CatB, ccMA cycloisomerase. Abbreviations: AV, acetovanillone; AVP, 4-acetyl-2-methoxyphenylphosphate; VAA, vanilloyl acetic acid; ccMA, *cis*,*cis*-muconic acid. (B) Schematic representations of the NGC703 recombinant strain, which contains pSEVA*acv* and pTS093*vce*-*aroY*. (C) Conversion of 1.2 mM AV by NGC703(pSEVA*acv* + pTS093*vce*-*aroY*) cells during growth in MMx-3 medium containing 15 g/L glucose. The concentrations of AV (red), VAA (purple), vanillic acid (green), protocatechuic acid (mustard), ccMA (blue), and *cis*,*trans*-muconic acid (gray) were periodically measured by HPLC. The concentration of glucose (triangles) was measured by a glucose electrode. Cell growth (squares) was monitored by measuring the OD_600_. All experiments were performed in triplicate, and each value represents the averages ± standard deviations.

We constructed pSEVA*acv* (RO1600/ColE1 *ori*), which carries *acvABCDEF* under the control of the *lac* promoter, and introduced pSEVA*acv* into NGC7. When resting cells of NGC7(pSEVA*acv*) (OD_600_ = 10.0) were incubated with 200 µM AV, AV was converted into VAA (Fig. S11A and B). In our previous study, we found VceA, which converts VAA and SAA into vanilloyl-CoA and syringoyl-CoA, respectively, and VceB, which converts vanilloyl-CoA and syringoyl-CoA into vanillic acid and syringic acid, respectively, from SYK-6, and that VAA can be converted into vanillic acid by these enzymes (Fig. 7A) (5). Thus, we constructed pTS093*vce* (RK2 *ori*), which carries *vceA* and *vceB* under the control of the *lac* promoter, and introduced pTS093*vce* into NGC7(pSEVA*acv*). When resting cells of NGC7(pSEVA*acv* + pTS093*vce*) (OD_600_ = 10.0) were incubated with 200 µM AV, AV was converted to vanillic acid (Fig. S11C). To produce ccMA from AV, we constructed pTS093*vce-aroY* (RK2 *ori*), which carries *aroY* with *vceA* and *vceB* under the control of the *lac* promoter, and then introduced pTS093*vce-aroY* into NGC703 with pSEVA*acv* (Fig. 7B). When the resulting strain was grown on an MMx-3 medium [34.2 g/L Na_2_HPO_4_×12H_2_O, 6.0 g/L KH_2_PO_4_, 1.0 g/L NaCl, 2.5 g/L (NH_4_)_2_SO_4_, 490 mg/L MgSO_4_×7H_2_O, 14.7 mg/L CaCl_2_×2H_2_O, and 5 mg/L FeSO_4_×7H_2_O] containing 15 g/L glucose as a carbon source, 1.2 mM AV could be converted into ccMA with 96% yield (mol ccMA/mol AV) (Fig. 7C). These results demonstrate that combining *acvABCDEF* with *vceA* and *vceB* is useful for the value-added chemical production from AV, a major aromatic monomer produced in oxidative/base-catalyzed lignin depolymerization.

## DISCUSSION

AV and AS are major lignin-derived aromatic monomers produced in alkaline oxidative depolymerization (13–16) and base-catalyzed depolymerization of lignin (17-19, 62-68). They are also contained in the black liquor produced during the kraft pulping process (20). This study identified *acvABCDEF* that encodes AV and AS carboxylase components and showed that these genes help produce value-added chemicals from AV.

As phenylphosphate synthases, Proteins 1 and 2 of *T. aromatica* K172 (51), PpsAB_GM_ of *Geobacter metallireducens* ATCC 53774 (69), and PpsAB_FP_ of *Ferroglobus placidus* DSM 10642 (70) have been reported. These enzymes are responsible for producing a phosphorylated intermediate essential for phenol carboxylation. Although not an example of carboxylation, 4-methylbenzyl phosphate synthase (CreHI) from *Corynebacterium glutamicum* produces a phosphorylated intermediate essential for oxidizing the 4-cresol methyl group (71). Based on the amino acid sequence similarity, AcvA corresponds to Protein 1, PpsA_GM_, PpsA_FP_, and CreH (40%–44% amino acid sequence identity), while AcvB corresponds to Protein 2, PpsB_GM_, PpsB_FP_, and CreI (39%–42% amino acid sequence identity) (Table S2). Among these phenylphosphate synthases, the reaction mechanism has been proposed in Proteins 1 and 2 of *T. aromatica* K172 (52). The phenol phosphorylation by Proteins 1 and 2 is similar to the ping-pong mechanism proposed for phosphoenolpyruvate synthase and is thought to proceed by the following two-step reactions. i) Protein 2 transfers the phosphoryl group from MgATP to His569 of Protein 1, producing His569-β-phosphate-γ-phosphate. ii) The terminal γ-phosphate is irreversibly hydrolyzed from Protein 1, and then the phosphate group of His569-β-phosphate is transferred to phenol to form phenylphosphate. AcvB, Protein 2, PpsB_GM_, PpsB_FP_, and CreI have an ATP binding domain (PPDK_N; PF01326) (Fig. S12). Additionally, AcvA, PpsA_GM_, PpsA_FP_, and CreH contained a His residue corresponding to Protein 1 His569 (Fig. S13). These facts suggest that AcvAB, PpsAB_GM_, PpsAB_FP_, and CreHI phosphorylate the hydroxy group of each substrate by a mechanism common to Proteins 1 and 2 of *T. aromatica* K172.

AcvF dephosphorylated AVP and ASP produced from AV and AS, respectively. In *T. aromatica* K172, phenylphosphate produced by Proteins 1 and 2 is dephosphorylated by the δ subunit of phenylphosphate carboxylase or non-enzymatic reaction generating phenolate anion. This intermediate is used as a substrate for carboxylation by the core enzyme [(αβγ)_3_] of phenylphosphate carboxylase (57, 58). Although AcvF and the 8 subunit of phenylphosphate carboxylase have no significant similarity with each other (7% identity), both AcvF and the δ subunit are classified in the HAD-like superfamily. Relatively low overall sequence similarity has been reported for this superfamily of enzymes (72). Therefore, AcvF probably dephosphorylates AVP/ASP, similar to the phenylphosphate carboxylase δ subunit, producing an anionic intermediate for AV/AS.

Crude AcvCDE converted AVP into VAA (Fig. S10C and D). This result suggests that phosphatase(s) present in UT26S complemented the AcvF function. Since *acvF* is essential for SYK-6 growth on AV/AS, the dephosphorylation of AVP/ASP catalyzed by AcvF appears crucial for AV/AS carboxylation. To verify this, AcvCDE purification and functional analysis using the purified enzyme are necessary for the future.

AcvC, AcvD, and AcvE were predicted to be BCCP, BC, and CT, respectively, based on the amino acid sequence similarity. In addition to these components, biotin-dependent carboxylases require biotin-protein ligase (BPL), which specifically adds biotin to a lysine residue of BCCP (73). A search for putative BPL genes in SYK-6 revealed the presence of SLG_23040, which showed 23% amino acid sequence identity with BirA of *B. subtilis* 168 (74). AVP was converted to VAA via *acvCDE* expression in *S. japonicum* UT26S, suggesting that AcvC (BCCP) was biotinylated by BPL present in UT26S. SJA_C1-13370, which showed 44% amino acid sequence identity with SLG_23040, may have functioned as BPL. *xccA*, *xccC*, and *xccB*, encoding putative biotin-dependent carboxylase components of *A. aromaticum* EbN1, are thought to be involved in 4’-hydroxyacetophenone carboxylation, which is structurally similar to AV (55). Additionally, *apkA*-*apkB*-*apkC*, encoding putative biotin-dependent carboxylase components of *R. rhodochrous* GD02, were recently predicted to be involved in AV catabolism (20). However, these enzyme genes have not been functionally characterized. To the best of our knowledge, this is the first report demonstrating that biotin-dependent carboxylase is involved in the catabolism of aromatic compounds. Phenol carboxylase of *T. aromatica* K172 (50-52, 57-59, 75) and acetophenone carboxylase [Apc(αα’βγ)_2_ core complex and Apcε] of *A. aromaticum* EbN1 (76, 77) are not biotin-dependent enzymes and show no similarity to AcvC, AcvD, and AcvE. Altogether, AV/AS carboxylation in SYK-6 seems to proceed as follows (Fig. S14). i) AV/AS is converted into AVP/ASP by transferring the phosphate group from MgATP to the hydroxy group of AV/AS by AcvAB. ii) AcvF dephosphorylates the resulting AVP/ASP to produce an anionic intermediate. iii) The anionic intermediate is converted to VAA/SAA by transferring the carboxyl group from the carboxylated biotin by AcvCDE.

In SYK-6, AV/AS is converted to VAA/SAA, an intermediate metabolite in the β-aryl ether catabolism (Fig. 1). Therefore, we investigated whether β-aryl ether catabolic genes and AV/AS catabolic genes coexist in the genomes of *Altererythrobacter* sp. strain Root672, *Altererythrobacter atlanticus* 26DY36, *Erythrobacter* sp. strain SG61- 1L, *Sphingobium* sp. strain 66-54, and *Sphingobium* sp. strain B12D2A, which have orthologs of the β-aryl ether catabolic genes (Table S3). Orthologs of *acvABCDEF* showing more than 45% amino acid sequence identity were found in all strains.

Particularly, the gene order of *acvABCDEF* in *Altererythrobacter atlanticus* 26DY36, *Sphingobium* sp. 66-54, and *Sphingobium* sp. B12D2A was conserved. Therefore, these strains may have evolved an AV/AS catabolic pathway by connecting the AV/AS carboxylase and the downstream pathway enzymes of the β-aryl ether catabolism.

Finally, we successfully achieved the microbial AV conversion (1.2 mM) into ccMA with 96% yield (mol ccMA/mol AV) using *Pseudomonas* sp. NGC703(pSEVA*acv* + pTS093*vce*-*aroY*), which is a *pcaHG catB* NGC7 mutant carrying *acvABCDEF*, *vceA*, *vceB*, and *aroY* (Fig. 7) (5, 34). In producing value-added chemicals from lignin through biological funneling, it is necessary to convert all degradation products obtained by chemical depolymerization of lignin or to degrade compounds that cannot be converted into products and not leave them in the culture medium to facilitate product purification. The results of this study will provide invaluable insights for improving the yield and purity of products in the biological conversion of aromatics obtained by alkaline oxidative and base-catalyzed depolymerization of lignin.

## MATERIALS AND METHODS

### Bacterial strains, plasmids, and culture conditions

The strains and plasmids used in this study are listed in Table 1. *Sphingobium* sp. strain SYK-6 and its mutants were grown in lysogeny broth (LB), Wx-SEMP, and Wx-SEMP containing 5 mM AV at 30°C. *Sphingobium japonicum* UT26S, *Pseudomonas* sp. NGC7, and its mutants were grown in LB at 30°C. When necessary, 12.5 mg/L nalidixic acid, 100 mg/L streptomycin, 25–50 mg/L kanamycin, or 12.5–15.0 mg/L tetracycline was added to the cultures. *Escherichia coli* strains were grown in LB at 37°C. For cultures of cells carrying antibiotic resistance markers, the media for *E. coli* transformants were supplemented with 100 mg/L ampicillin, 25 mg/L kanamycin, or 12.5 mg/L tetracycline.

**TABLE 1.** Strains and plasmids used in this study.

| Strain or plasmid | Relevant characteristic(s)<sup>a</sup> | Reference or source |
| --- | --- | --- |
| **Strains** |  |  |
| *Sphingobium* sp. |  |  |
| SYK-6 | Wild type; NBRC 103272/JCM 17495, NaI<sup>r</sup> Sm<sup>r</sup> | (84) |
| Δ06490 (SME062) | SYK-6 derivative; ΔSLG_06490 (*acvF*), NaI<sup>r</sup> Sm<sup>r</sup> | This study |
| Δ06500 (SME063) | SYK-6 derivative; ΔSLG_06500 (*acvE*), NaI<sup>r</sup> Sm<sup>r</sup> | This study |
| Δ06510 (SME064) | SYK-6 derivative; ΔSLG_06510 (*acvD*), NaI<sup>r</sup> Sm<sup>r</sup> | This study |
| Δ06520 (SME065) | SYK-6 derivative; ΔSLG_06520 (*acvC*), NaI<sup>r</sup> Sm<sup>r</sup> | This study |
| Δ06530 (SME066) | SYK-6 derivative; ΔSLG_06530 (*acvB*), NaI<sup>r</sup> Sm<sup>r</sup> | This study |
| Δ06540 (SME067) | SYK-6 derivative; ΔSLG_06540 (*acvA*), NaI<sup>r</sup> Sm<sup>r</sup> | This study |
| Δ06550 (SME068) | SYK-6 derivative; ΔSLG_06550 (*acvR*), NaI<sup>r</sup> Sm<sup>r</sup> | This study |
| *Sphingobium japonicum* |  |  |
| UT26S | Type strain, NBRC 101211/JCM17232, γ-hexachlorocyclohexane degradation | (85) |
| *Pseudomonas* sp. |  |  |
| NGC7 | Wild type | (34) |
| NGC703 | NGC7 derivative; Δ*pcaHG catB* | (34) |
| *Escherichia coli* |  |  |
| BL21(DE3) | F<sup>-</sup> *ompT hsdSB*(r<sub>B</sub><sup>-</sup> m<sub>B</sub><sup>-</sup>) *gal dcm* (DE3); T7 RNA polymerase gene under the control of the *lacUV5* promoter | (86) |
| HB101 | *recA13 supE44 hsd20 ara-14 proA2 lacY1 galK2 rpsL20 xyl-5 mtl-1* | (87) |
| NEB 10-beta | Δ(*ara-leu*) 7697 *araD139 fhuA ΔlacX74 galK16 galE15 e14- φ80dlacZΔM15 recA1 relA1 endA1 nupG rpsL* (Sm<sup>r</sup>) *rph spoT1 Δ(mrr-hsdRMS-mcrBC)* | New England Biolabs |
| **Plasmids** |  |  |
| pRK2013 | Tra<sup>+</sup> Mob<sup>+</sup> ColE1 replicon, Km<sup>r</sup> | (88) |
| pAK405 | Plasmid for allelic exchange and markerless gene deletions in sphingomonads, Km<sup>r</sup> | (82) |
| pJB861 | RK2 broad-host-range expression vector; P<sub>m</sub> *xylS*, Km<sup>r</sup> | (89) |
| pET-21a(+) | Expression vector; T7 promoter, Ap<sup>r</sup> | Novagen |
| pET-16b | Expression vector; T7 promoter, Ap<sup>r</sup> | Novagen |
| pQF | Expression vector; Q5 promoter, codon-optimized *cymR*, Tc<sup>r</sup> | (90) |
| pUC118 | Cloning vector; lactose promoter (P<sub>lac</sub>), Ap<sup>r</sup> | Takara Bio |
| pSEVA241 | pRO1600/ColE1 replicon, Km<sup>r</sup> | (83) |
| pTS093 | pJB866 with a 0.2-kb PCR amplified fragment containing P<sub>lac</sub> from pUC118 | (12) |
| pTS032 | pUC118 with a 1.6-kb KpnI fragment carrying *aroY* | (35) |
| pJHV01 | pJB866 with a 4.6-kb BamHI PCR amplified fragment carrying *hvpZ*, SLG_20400, *vceA*, and *vceB* | (5) |
| pAK06490 | pAK405 with a 1.7-kb deletion cassette carrying up-and downstream regions of *acvF* | This study |
| pAK06500 | pAK405 with a 1.6-kb deletion cassette carrying up-and downstream regions of *acvE* | This study |
| pAK06510 | pAK405 with a 1.6-kb deletion cassette carrying up-and downstream regions of *acvD* | This study |
| pAK06520 | pAK405 with a 1.4-kb deletion cassette carrying up-and downstream regions of *acvC* | This study |
| pAK06530 | pAK405 with a 1.6-kb deletion cassette carrying up-and downstream regions of *acvB* | This study |
| pAK06540 | pAK405 with a 1.7-kb deletion cassette carrying up-and downstream regions of *acvA* | This study |
| pAK06550 | pAK405 with a 1.8-kb deletion cassette carrying up-and downstream regions of *acvR* | This study |
| pE21*acv* | pET-21a(+) with a 7.4-kb PCR amplified fragment carrying *acvABCDEF* | This study |
| pJB*acv* | pJB861 with a 7.4-kb NotI fragment carrying *acvABCDEF* from pE21*acv* | This study |
| pE16*acvA* | pET-16b with a 1.8-kb PCR amplified fragment carrying *acvA* | This study |
| pE16*acvB* | pET-16b with a 1.0-kb PCR amplified fragment carrying *acvB* | This study |
| pE16*acvF* | pET-16b with a 0.8-kb PCR amplified fragment carrying *acvF* | This study |
| pE16*acvCDE* | pET-16b with a 3.4-kb PCR amplified fragment carrying *acvCDE* | This study |
| pQF*acvCDE* | pQF with a 3.3-kb PCR amplified fragment carrying *acvCDE* | This study |
| pSEVA241_P<sub>lac</sub> | pSEVA241 with a 0.2-kb PCR amplified fragment containing P<sub>lac</sub> from pUC118 | This study |
| pSEVA*acv* | pSEVA241_P<sub>lac</sub> with a 7.4-kb NotI fragment carrying *acvABCDEF* from pJB*acv* | This study |
| pQF*vce* | pQF with a 1.4-kb PCR amplified fragment carrying *vceA* and *vceB* from pJHV01 | This study |
| pTS093*vce* | pTS093 with a 1.4-kb PCR amplified fragment carrying *vceA* and *vceB* from pQF*vce* | This study |
| pTS093*vce-aroY* | pTS093*vce* with a 1.6-kb KpnI fragment carrying *aroY* from pTS032 | This study |

<sup>a</sup>NaI<sup>r</sup>, Sm<sup>r</sup>, Km<sup>r</sup>, Tc<sup>r</sup>, and Ap<sup>r</sup>, resistance to nalidixic acid, streptomycin, kanamycin, tetracycline, and ampicillin, respectively.

### Preparation of substrates

VAA and SAA were prepared as described previously (5). For the preparation of AVP and ASP, AV or AS (final concentration: 200 µM) was incubated in 1 mL 50 mM Tris-HCl buffer (pH 7.5; Buffer A) containing the cell extracts of *E. coli* BL21(DE3) cells harboring pE16*acvA* and *E. coli* BL21(DE3) cells harboring pE16*acvB* (1 mg protein/mL each), 2 mM ATP, 2 mM MgCl_2_, and 200 μM MnCl_2_. After incubation for 2 h at 30°C, complete consumption of AV or AS was confirmed by HPLC, and then the reaction mixtures were filtered using an Amicon Ultra spin filter unit (10 kDa cutoff; Millipore). *Cis*,*trans*-muconic acid was prepared by incubating ccMA (1.0 g/L) for 2 h at 60°C in a solution whose pH was adjusted to 4.0 with acetic acid. After the incubation, the pH was adjusted to 7 with sodium hydroxide and it was stored at -80°C until use. Other aromatic compounds were purchased from Tokyo Chemical Ind., Co., Ltd.; Sigma-Aldrich Co., LLC.; and FUJIFILM Wako Pure Chemical Corporation.

### Identification of the metabolites

SYK-6 cells grown in LB were inoculated into Wx-SEMP to an OD_600_ of 0.2 and grown at 30°C. AV (5 mM) was added when the OD_600_ of the culture reached 0.5, and the culture was then further incubated for 12 h. Cells were collected by centrifugation (5,000 × *g* for 5 min at 4°C), washed twice with Wx minimal medium, and resuspended in the same medium. The resultant cell suspensions were inoculated into Wx medium containing 1 mM AV to an OD_600_ of 0.2 and incubated for 33 h at 30°C. The reaction mixtures were centrifuged, and the supernatants were collected. The resulting filtered samples were analyzed by HPLC– MS.

### HPLC–MS analysis

HPLC–MS analysis was performed with the Acquity UPLC system coupled with an Acquity TQ detector as described previously (78). The in vivo reaction products of AV and AS were analyzed using a UPLC equipped with an Acquity UPLC BEH C18 column (2.1 by 100 mm; Waters). The flow rate of the mobile phase was 0.5 mL/min. The in vitro reaction products of AV, AS, acetophenone, 4’- hydroxyacetophenone, 3’-hydroxyacetophenone, 3’,4’-dihydroxyacetophenone, 3’- hydroxy-4’-methoxyacetophenone, 3’,4’-dimethoxyacetophenone, 3’,4’,5’- trimethoxyacetophenone, 4’-hydroxypropiophenone, 4’-hydroxybuthyrophenone, 4’- hydroxyvalerophenone, guaiacol, vanillic acid, 4-hydroxybenzoic acid, AVP, and ASP were analyzed using a UPLC equipped with a TSKgel ODS-140HTP column (2.1 by 100 mm; Tosoh). The flow rate of the mobile phase was 0.5 mL/min. The mobile phase was a mixture of Solution A (acetonitrile containing 0.1% formic acid) and Solution B (water containing 0.1% formic acid) with the following conditions: *(i) detection of in vivo reaction products of AV*: 0 to 3.0 min, linear gradient from 5 to 15% A; 3.0 to 4.0 min, decreasing gradient from 15 to 5% A. *(ii) detection of in vivo reaction products of AS*: 0 to 3.5 min, 10% A; 3.5 to 4.0 min, linear gradient from 10 to 30% A; 4.0 to 5.0 min, 30% A; 5.0 to 5.1 min, decreasing gradient from 30 to 10% A; 5.1 to 6.0 min, 10% A. *(iii) detection of the in vitro reaction products*: 0 to 0.5 min, 2% A; 0.5 to 1.5 min, linear gradient from 2 to 15% A; 1.5 to 3.0 min, linear gradient from 15 to 100% A; 3.0 to 4.0 min, decreasing gradient from 100 to 2% A; 4.0 to 5.0 min, 2% A. AV, AS, acetophenone, 4’-hydroxyacetophenone, 3’-hydroxyacetophenone, 3’,4’- dihydroxyacetophenone, 3’-hydroxy-4’-methoxyacetophenone, 3’,4’- dimethoxyacetophenone, 3’,4’,5’-trimethoxyacetophenone, 4’-hydroxypropiophenone, 4’-hydroxybuthyrophenone, 4’-hydroxyvalerophenone, guaiacol, vanillic acid, 4- hydroxybenzoic acid, AVP, ASP, VAA, and SAA were detected at 275, 300, 255, 275, 260, 275, 274, 274, 280, 271, 272, 271, 276, 260, 255, 303, 300, 280, and 308 nm, respectively. In the ESI-MS analysis, MS spectra were obtained using the negative-ion and positive-ion modes with the settings reported in our previous study (78).

### Resting cell assay

SYK-6 cells grown in LB were inoculated into Wx-SEMP to an OD_600_ of 0.2 and grown at 30°C until the OD_600_ of the culture reached 0.5. Following the addition of 5 mM AV, the cells were incubated for a further 12 h as the inducing condition. For the non-inducing condition, the culture was incubated for a further 12 h without AV. The cells were collected by centrifugation (5,000 × *g* for 5 min at 4°C) and then washed twice with Buffer A. The cells were resuspended in the same buffer and used as resting cells. Preparation of resting cells of *S. japonicum* UT26S are described below.

Resting cells of SYK-6 (OD_600_ of 5.0) were incubated with 1.0 mM AV at 30°C with shaking. Resting cells of UT26S harboring pJB861and UT26S harboring pJB*acv* (OD_600_ of 10.0) were incubated with 200 µM AV or AS at 30°C with shaking. Portions of the reaction mixtures were collected, and the amounts of compounds were measured by HPLC.

### Analysis of nucleotide and amino acid sequences

Nucleotide sequences were determined by Eurofins Genomics. Sequence analysis was performed using the MacVector program (MacVector, Inc.). Sequence similarity searches, pairwise alignments, and multiple alignments were conducted using the BLASTP program (79), the EMBOSS Needle program through the EMBL-EBI server (80), and the Clustal Omega program (81), respectively.

### RT-PCR analysis

SYK-6 cells grown in LB were inoculated into Wx-SEMP to an OD_600_ of 0.2 and grown at 30°C. AV (5 mM) was added when the OD_600_ of the culture reached 0.5, and the culture was then further incubated for 6 h. Total RNA was isolated from the cells using an Illumina RNAspin Mini RNA isolation kit (GE Healthcare). The samples were treated with DNase I to remove any contaminating genomic DNA. Total RNA (4 μg) was reverse transcribed using SuperScript IV reverse transcriptase (Invitrogen) with random hexamer primers. The cDNA was purified using a NucleoSpin Gel and PCR Clean-up kit (Takara Bio, Inc.). PCR was performed with the cDNA, specific primers (Table 2), and Q5 High-Fidelity DNA Polymerase (New England Biolabs). The DNA obtained was electrophoresed on a 0.8% agarose gel.

**TABLE 2.**
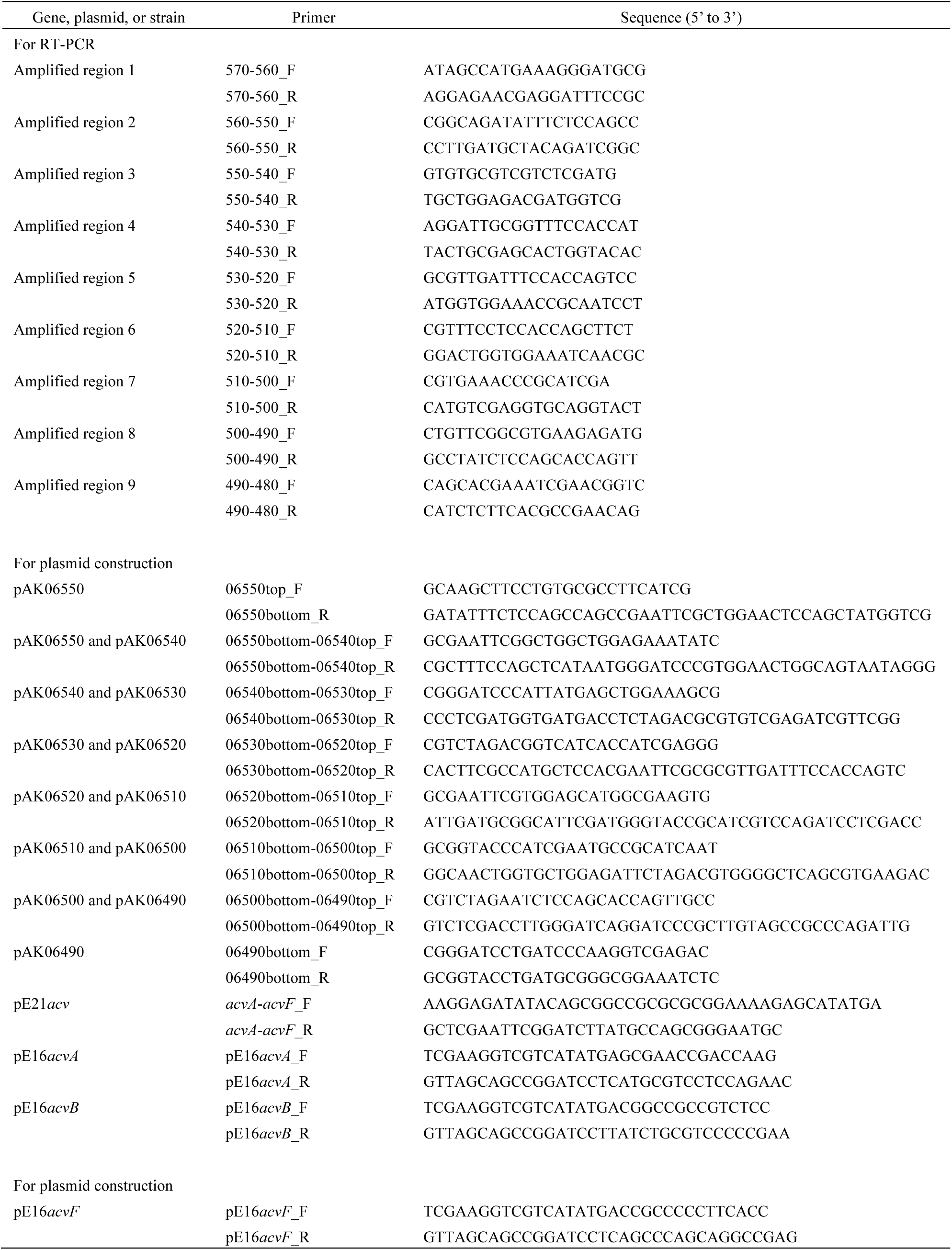
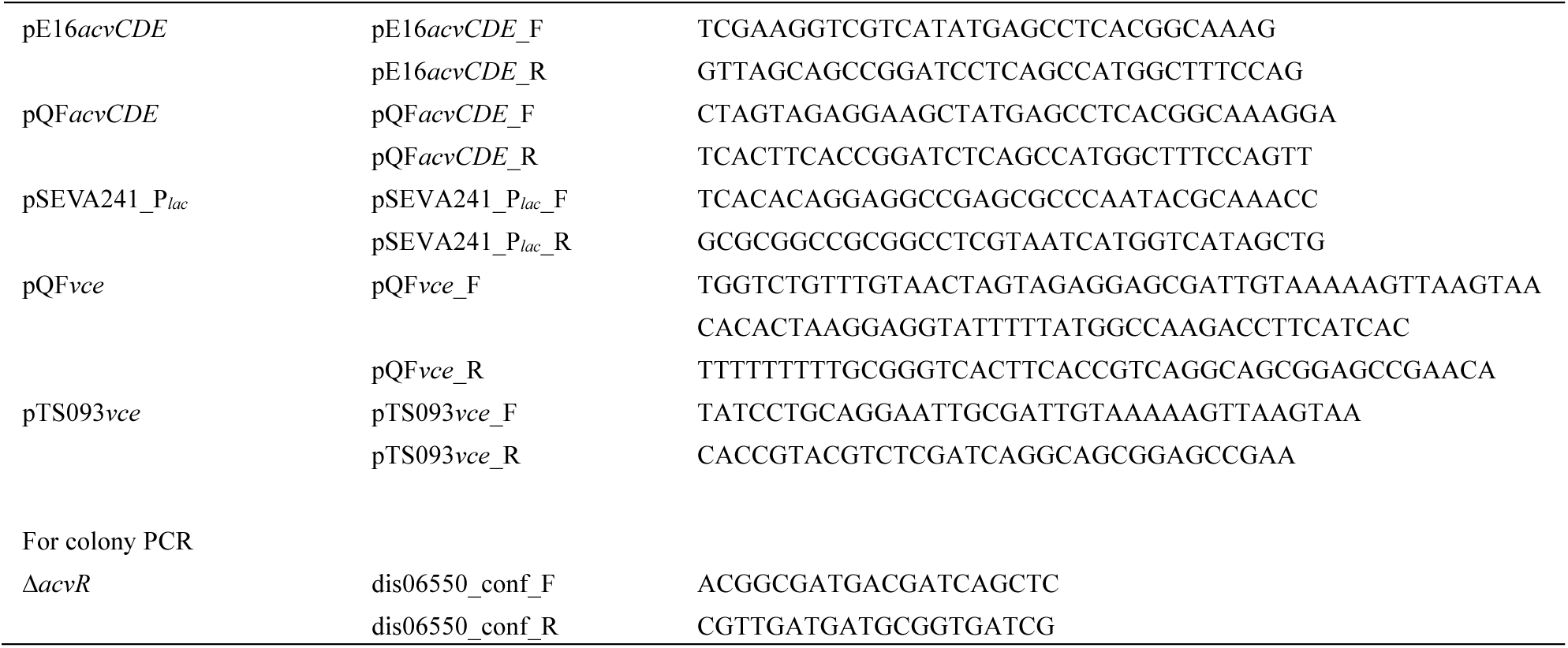
Primers used in this study.

### Construction of mutants

To construct the mutants, the upstream and downstream regions (0.7−1.0 kb each) of the genes were amplified by PCR from SYK-6 total DNA using the primer pairs listed in Table 2. The amplified fragments were ligated into pAK405 (Kaczmarczyk et al., 2012). Each of the resulting plasmids was introduced into SYK-6 cells by triparental mating, and the resulting mutants were selected as described previously (82). Disruption of each gene was examined by Southern hybridization analysis using the digoxigenin system (Roche) or colony PCR using the primer pairs listed in Table 2.

### Growth measurement

SYK-6 and its mutant cells grown in LB were inoculated into Wx-SEMP to an OD_600_ of 0.2 and grown at 30°C. AV (5 mM) was added when the OD_600_ of the culture reached 0.5, and the culture was then further incubated for 12 h.

Cells were collected by centrifugation (5,000 × *g* for 5 min at 4°C), washed twice with Wx minimal medium, and resuspended in the same medium. The resultant cell suspensions were inoculated into Wx medium containing 1 mM AV or AS to an OD_660_ of 0.2. Cells were incubated at 30°C with shaking (60 rpm) and cell growth was monitored every hour by measuring the OD_660_ with a TVS062CA biophotorecorder (Advantec Co., Ltd.). After 52 h (AV) and 48 h (AS) cultivation, 1 mM AV or AS was added to the culture medium.

### Expressions of *acvABCDEF* in heterologous hosts and enzyme purification

For expression of *acvABCDEF* in *S. japonicum* UT26S, a 7.4-kb fragment carrying *acvABCDEF* with the NotI site at 5’ terminus and 3’ terminus was amplified by PCR. The amplified fragment was cloned into the BamHI site of pET-21a(+) using In-Fusion HD cloning Kit (Takara Bio), and the NotI fragment of the resulting plasmid was then inserted in pJB861 to generate pJB*acv*. For expression of *acvA*, *acvB*, *acvF*, and *acvCDE* in *E. coli*, DNA fragments carrying each gene were amplified by PCR from the SYK-6 total DNA using primer pairs shown in Table 2. The amplified fragments were cloned into NdeI and BamHI sites of pET-16b using In-Fusion HD cloning Kit. For expression of *acvCDE* in UT26S, DNA fragment carrying *acvCDE* was amplified by PCR from the SYK-6 total DNA (Table 2). The amplified fragment was cloned into BamHI site of pQF using an NEBuilder HiFi DNA assembly cloning kit (New England Biolabs) to generate pQF*acvCDE*. Nucleotide sequences of the resultant plasmids were then confirmed.

pJB*acv* and pQF*acvCDE* were introduced into UT26S cells by electroporation. Cells of UT26S harboring pJB*acv* were inoculated into LB supplemented with 1 mM *m*-toluic acid as an inducer and grown at 30°C for 24 h. Cells of UT26S harboring pQF*acvCDE* were inoculated into LB supplemented with 0.1 mM 4-isopropylbenzoic acid as an inducer and grown at 30°C for 24 h. Cells of *E. coli* BL21(DE3) harboring pE16*acvA*, pE16*acvB*, pE16*acvF*, or pE16*acvCDE* were grown in LB at 30°C. Each gene expression was induced for 4 h at 30°C by adding 1 mM isopropyl-β-D-thiogalactopyranoside when the OD_600_ of the culture reached 0.5. The cells of UT26S and *E. coli* transformants were then collected by centrifugation (5,000 × *g* for 5 min at 4°C), washed twice with buffer A, resuspended in the same buffer, and used as resting cells. The cells were then disrupted using an ultrasonic disintegrator. After centrifugation (19,000 × *g* for 15 min at 4°C), the supernatants were obtained as cell extracts (crude enzymes). AcvA, AcvB, and AcvF were purified from cell extracts of *E. coli*(pE16*acvA*), *E. coli*(pE16*acvB*), and *E. coli*(pE16*acvF*), respectively, using a His SpinTrap column (GE Healthcare). Resultant elution fractions were subjected to desalting and concentration using an Amicon Ultra centrifugal filter unit (30 kDa cutoff; Merck Millipore), and the enzyme preparations were stored at -80°C. SDS-PAGE and protein concentration determination using the Bradford method were performed as described previously (43).

### Enzyme activity of AcvAB

Crude AcvA, crude AcvB, crude AcvA + AcvB (500 µg protein/mL each) or purified AcvA + AcvB (100 µg protein/mL each) were incubated with 100 or 200 µM AV in the presence of 2 mM ATP, 2 mM MgCl_2_, and 200 µM MnCl_2_ in buffer A for 10–30 min at 30°C. The reactions were stopped by adding acetonitrile to a final concentration of 50%. Protein precipitates were removed through centrifugation (19,000 × *g* for 15 min). The resulting supernatants were diluted with water to a final concentration of acetonitrile of 25%, filtered, and analyzed using HPLC–MS. Specific activity was expressed in moles of AV converted per minute per milligram of protein.

### Enzyme properties of AcvAB

The enzyme reaction was conducted by incubating crude AcvA + AcvB (10–1000 µg protein/mL each) with 200 µM AV, 2 mM ATP, 2 mM MgCl_2_, and 200 µM MnCl_2_ in buffer A for 10 min at 30°C. After incubation, the amounts of AV were measured using HPLC. To examine cofactor requirement of AcvAB, crude AcvA + AcvB (50–1000 µg protein/mL each) was incubated with 100 µM AV in the presence and absence of cofactors (2 mM ATP, 2 mM MgCl_2_, 200 µM MnCl_2_, 2 mM ATP + 2 mM MgCl_2_, 2 mM ATP + 200 µM MnCl_2_, 2 mM MgCl_2_ + 200 µM MnCl_2_, and 2 mM ATP + 2 mM MgCl_2_ + 200 µM MnCl_2_) for 10 min at 30°C. To determine the substrate range, 100 µM AV, AS, acetophenone, 4’- hydroxyacetophenone, 3’-hydroxyacetophenone, 3’,4’-dihydroxyacetophenone, 3’- hydroxy-4’-methoxyacetophenone, 3’,4’-dimethoxyacetophenone, 3’,4’,5’- trimethoxyacetophenone, 4’-hydroxypropiophenone, 4’-hydroxybuthyrophenone, 4’- hydroxyvalerophenone, guaiacol, vanillic acid, and 4-hydroxybenzoic acid were used for the reaction, and the conversion of substrates and generation of reaction products were analyzed using HPLC–MS.

### Enzyme activity of AcvF

Purified AcvF (5 µg protein/mL) was incubated with 100 µM AVP or ASP in buffer A for 60 min at 30°C. The supernatants were then analyzed using HPLC.

### Enzyme activity of AcvCDE

Cell extract of *E. coli* BL21(DE3) harboring pE16*acvCDE*, cell extract of *S. japonicum* UT26S harboring pQF, or cell extract of UT26S harboring pQF*acvCDE* (1 mg protein/mL) was incubated with 100 µM AV or AS, 2 mM ATP, 2 mM MgCl_2_, 200 µM MnCl_2_, and 10 mM NaHCO_3_ in the presence or absence of crude AcvA + AcvB (1 mg protein/mL each) and purified AcvF (10 µg protein/mL) in buffer A for 60 min at 30°C. When using AVP as a substrate, cell extract of UT26S harboring pQF or cell extract of UT26S harboring pQF*acvCDE* (1 mg protein/mL) was incubated with 100 µM AVP, 2 mM ATP, 2 mM MgCl_2_, 200 µM MnCl_2_, and 10 mM NaHCO_3_ in buffer A for 60 min at 30°C. The supernatants were then analyzed using HPLC.

### ccMA production from AV

A 0.2-kb fragment carrying the *lac* promoter (P*_lac_*) from pUC118 was amplified and cloned into the SfiI site of pSEVA241 (83) with In- Fusion HD cloning kit to generate pSEVA241_P*_lac_*. The 7.2-kb NotI fragment carrying *acvABCDEF* from pJB*acv* was ligated into the corresponding site of pSEVA241_P*_lac_* to generate pSEVA*acv*. A 1.5-kb fragment carrying *vceA* and *vceB* was amplified by PCR using pJHV01 (5) and the primer pairs listed in Table 2. The resulting fragment was cloned into the BamHI site of pQF by NEBuilder HiFi DNA assembly cloning kit to generate pQF*vce*. The 1.5-kb fragment carrying *vceA* and *vceB* was amplified by PCR using pQF*vce* and the primer pairs listed in Table 2. The resulting fragment was cloned into the EcoRI-XhoI site of pTS093 (12) with In-Fusion HD cloning kit to generate pTS093*vce*. The 1.6-kb KpnI fragment carrying *aroY* from pTS032 (35) was ligated into the corresponding sites of pTS093*vce* to generate pTS093*vce*-*aroY*. pSEVA241_P*_lac_*, pSEVA*acv*, or pSEVA*acv* + pTS093*vce* was introduced into NGC7 cells by electroporation. pSEVA*acv* and pTS093*vce*-*aroY* were introduced into NGC703 (ϕ..*pcaHG catB*) cells by electroporation.

### Conversion of AV by NGC7 transformants

Cells of NGC7 harboring pSEVA241_P*_lac_*, NGC7 harboring pSEVA*acv*, and NGC7 harboring pSEVA*acv* + pTS093*vce* were grown in LB containing kanamycin or kanamycin + tetracycline for 16 h. The cells were collected by centrifugation at 9,000 × *g* for 3 min, washed twice with MMx-3 buffer [34.2 g/L Na_2_HPO_4_×12H_2_O, 6.0 g/L KH_2_PO_4_, 1.0 g/L NaCl, and 2.5 g/L (NH_4_)_2_SO_4_], resuspended in the same buffer, and used as resting cells. Resting cells of NGC7 harboring pSEVA241_P*_lac_*, NGC7 harboring pSEVA*acv*, and NGC7 harboring pSEVA*acv* + pTS093*vce* (OD_600_ of 10.0) were incubated with 200 µM AV or AS at 30°C with shaking. Portions of the reaction mixtures were periodically collected, and the reactions were stopped by centrifugation. The resultant supernatants were diluted, filtered, and analyzed using an HPLC instrument 1200 series (Agilent Technologies Inc.) equipped with a ZORBAX Eclipse Plus C18 column (4.6 by 150 mm; Agilent Technologies Inc.). The flow rate of the mobile phase was 1.0 mL/min, and the detection wavelength was 280 nm. The mobile phase was a mixture of Solution A (50% methanol containing 1% acetic acid) and Solution B (5% methanol containing 1% acetic acid) with the following conditions: 0–8.0 min, linear gradient from 0% to 20% A; 8.0– 13.0 min, linear gradient from 20% to 100% A; 13.0–18.0 min, 100% A; 18.0–23.0 min, decreasing gradient from 100% to 0% A; 23.0–25.5 min, 100% B.

Cells of NGC703(pSEVA*acv* + pTS093*vce-aroY*) were grown in LB containing kanamycin and tetracycline for 16 h. The cells were collected by centrifugation at 9,000 × *g* for 3 min, washed twice with MMx-3 medium, and resuspended in 5 mL of the same medium. The cells were then inoculated into 10 mL of MMx-3 medium containing 15 g/L glucose, 1.2 mM AV, kanamycin, and tetracycline to an OD_600_ of 0.1 and incubated with shaking for 70 h at 30°C. Cell growth was measured by OD_600_.

Portions of the cultures were periodically collected, and the reactions were stopped by centrifugation. The resultant supernatants were diluted, filtered, and analyzed using a HPLC instrument 1200 series. ccMA yields were calculated as [the produced ccMA (mol)/the consumed AV (mol)] × 100%. The concentrations of glucose in the culture were measured with a BF-5i biosensor (Oji Scientific Instruments, Ltd.).

## ACKNOWLEDGEMENT

This work was supported by JST-Mirai Program Grant Number JPMJMI19E2, Japan, a grant from the Ministry of Agriculture, Forestry and Fisheries of Japan (Rural Biomass Research Project BM-D1310), and JSPS KAKENHI Grant Number JP26850046.

